# Condensin’s ATPase Machinery Drives and Dampens Mitotic Chromosome Condensation

**DOI:** 10.1101/216630

**Authors:** Ahmed M.O. Elbatsh, Jonne A. Raaijmakers, Robin H. van der Weide, Jelmi Kuit de Bos, Hans Teunissen, Sol Bravo, René H. Medema, Elzo de Wit, Christian H. Haering, Benjamin D. Rowland

**Affiliations:** Division of Gene Regulation, the Netherlands Cancer Institute, Plesmanlaan 121, 1066 CX Amsterdam, the Netherlands; Division of Cell Biology, the Netherlands Cancer Institute, Plesmanlaan 121, 1066 CX Amsterdam, the Netherlands; Cell Biology and Biophysics Unit, Structural and Computational Biology Unit, European Molecular Biology Laboratory (EMBL), Heidelberg, Germany

## Abstract

Chromosome condensation by condensin is essential for faithful chromosome segregation. Metazoans have two complexes, named condensin I and II. Both are thought to act by creating looped structures in DNA, but how they do so is unknown. Condensin’s SMC subunits together form a composite ATPase with two pseudo-symmetric ATPase sites. We reveal that these sites have opposite functions in the condensation process. One site drives condensation, while the other site rather has a dampening function. Mutation of this dampener site hyperactivates both condensin I and II complexes. We find that hyperactive condensin I efficiently shortens chromosomes in the total absence of condensin II. The two complexes form loops with different lengths, and specifically condensin II is key to the decatenation of sister chromatids and the formation of a straight chromosomal axis.

## INTRODUCTION

Mitotic chromosome condensation is dependent on the condensin complex. Condensin is a ring-shaped structure that at its heart has a heterodimer of SMC2 and SMC4 proteins that together form a composite ABC-like ATPase. This heterodimer binds three non-SMC subunits, one known as the kleisin, which together with the SMCs forms a tri-partite ring, and two HEAT-repeat proteins (Hirano, 2016). Condensin can topologically entrap DNA inside its tripartite ring, and harbours a second DNA binding interface in one of its HEAT subunits (Cuylen et al., 2011; Kschonsak et al., 2017).

Metazoans express two condensin complexes called condensin I and condensin II (Hirota, 2004; Ono et al., 2003). The two complexes share the same SMC dimer, but each bind unique kleisin and HEAT subunits (Figure 1A). The complexes further differ in their cellular localization and residence time on chromatin (Gerlich et al., 2006; Shintomi and Hirano, 2011). Condensin II is always nuclear, whereas condensin I is cytoplasmic, and only gains access to chromosomes upon nuclear envelope breakdown (NEBD). In mitosis, condensin II binds to chromosomes with a longer residence time than condensin I.

**Figure 1.**
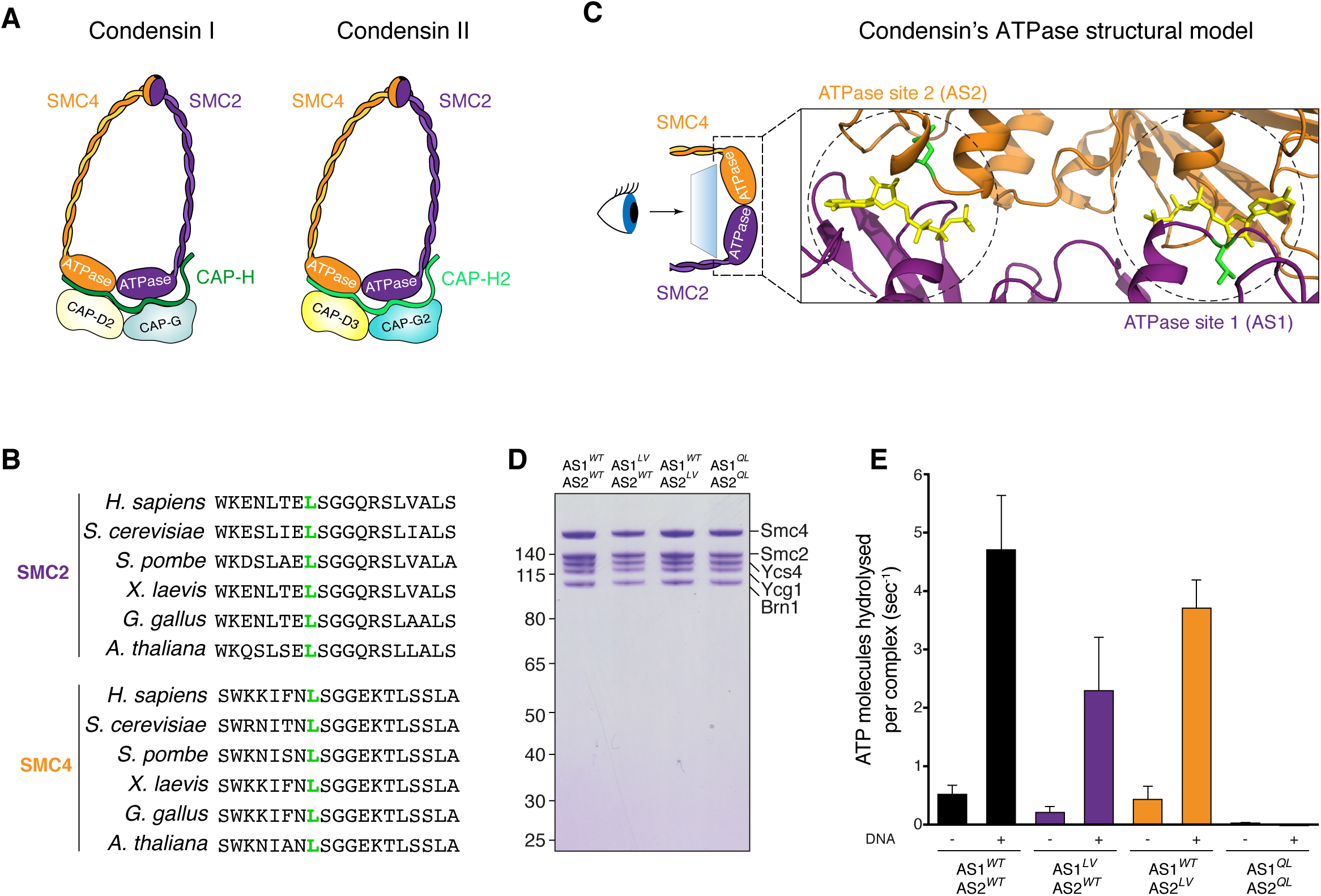
Condensin’s ATPase Sites Differentially Control its Hydrolysis Activity. (A) Schematic depiction of the condensin complex. Metazoans have two complexes, condensin I and II, which share the same SMC2-SMC4 heterodimer, but each has three complex-specific additional subunits. (B) Sequence alignment showing the conservation of a key amino acid in the signature motif of condensin’s SMC2 and SMC4 ATPase domains in all examined species. (C) Structural model of SMC2-SMC4 heterodimer showing the structural symmetry between the two proteins. The ATPase head domains of the two proteins engage by sandwiching two ATP molecules (shown in yellow). This yields two composite ATPase sites, which we call ATPase site 1 and 2 (AS1 and AS2, respectively). The model was generated using SWISS-MODEL, based on the crystal structures of cohesin’s subunits Smc1 (PDB: 1W1W) and Smc3 (PDB: 4UX3). SMC2 is shown in purple and SMC4 in orange. The SMC2 L1085 and SMC4 L1191 residues are highlighted in green. (D) Coomassie-stained SDS PAGE of the condensin holocomplex samples as used in (E). (E) ATP hydrolysis rates measured for condensin holocomplexes (5 μM) in the absence or presence of 10 μg/μl of a relaxed 6.4-kb plasmid DNA. Error bars represent SD of five independent experiments for wild-type, AS1^LV^ (SMC2 L1048V)-AS2^WT^ and AS1^WT^ – AS2^LV^ (SMC4 L1323V) mutants, and three independent experiments for the negative control using the Q-loop mutant AS1^QL^ (SMC2 Q147L)-AS1^QL^ (SMC4 Q302L).

Chromosome condensation has been suggested to involve the processive enlargement of chromatin loops along chromosomes by the concerted action of condensin I and condensin II (Goloborodko et al., 2016; Nasmyth, 2001). This ‘Loop extrusion’ model provides an explanation for the axial localization of condensin along mitotic chromosomes. It also explains why condensin does not accidentally create inter-chromosomal linkages. Recent experimental evidence indicates that the related cohesin complex and bacterial SMC act through this looping mechanism (Haarhuis et al., 2017; Wang et al., 2017). Loop extrusion therefore is likely to reflect a universal mechanism by which SMC complexes operate.

What drives mitotic chromosome condensation is an important unanswered question. A main suspect is condensin’s ATPase machinery. Condensin harbours two ATPase sites that each sandwich an ATP molecule between the signature motif of one SMC subunit, and the Walker A and Walker B motifs of the other. To investigate the role of this machinery in condensation, we made use of our recently identified ATPase mutants in the cohesin complex that affect its ATPase cycle, but do support viability (Elbatsh et al., 2016). These mutations substitute a universally conserved Leucine residue of the signature motif of either of cohesin’s ATPase sites by a Valine. As the ATPase machineries of cohesin and condensin are very similar, these residues are also conserved through the condensin complex (Figure 1B, S1A and S1B). We have used the analogous mutations in condensin to identify an asymmetric activity within condensin’s ATPase machinery, in which one ATPase site drives, and the second rather dampens the condensation process. We have also dissected the specific functions of the two condensin complexes.

## RESULTS

### Condensin’s ATPase Sites Differentially Control its Hydrolysis Activity

To investigate the role of condensin’s ATPase sites in chromosome condensation, we mutated each individual site. We will refer to the two ATPase sites as AS1 and AS2 respectively (Figure 1C). First we assessed the effects of the mutations on condensin’s ATPase activity. We purified the condensin holocomplex to homogeneity from budding yeast (Figure 1D), and performed ATPase assays using thin-layer chromatography and radiolabeled ATP (Figure 1E). Consistent with previous work, wild type condensin efficiently hydrolysed ATP, which was further stimulated by the addition of DNA. (Terekawa et al., 2017). Intriguingly, the AS1^LV^ and AS2^LV^ mutations affected hydrolysis to different degrees. Whereas AS1^LV^ mutation clearly impaired both the basic and the DNA-stimulated ATPase activity of the holocomplex, the AS2^LV^ mutation only mildly reduced its ATPase activity. Each ATPase site apparently has a distinct contribution to condensin’s ATPase activity.

### Condensin’s ATPase Sites Drive and Dampen Chromosome Condensation

Next, we studied the role of condensin’s ATPase sites in chromosome condensation. We mutated the endogenous allele of each individual ATPase site in the human HAP1 cell line using CRISPR/Cas9 genome-editing technology. We used guide RNAs that led to cleavage of either the *SMC2* or *SMC4* gene, and provided donor oligos that upon homology-directed repair introduced the desired mutations, and at the same time rendered the genes non-cleavable by the Cas9 nuclease (Figure S1C and S1D). We hereby successfully obtained HAP1 cells with mutant endogenous alleles of *SMC2* (AS1^LV^) and of *SMC4* (AS2^LV^).

We then prepared chromosome spreads from wild type and mutant cells, and scored for chromosome condensation. Surprisingly, each mutation led to a very different phenotype. The AS1^LV^ mutation resulted in major condensation defects. The chromosomes of these mutant cells were fuzzy, and the individual chromosomes were hard to discern (Figures 2A and 2B). By marked contrast, the AS2^LV^ mutation did not lead to condensation defects. Chromosomes from these mutant cells compacted well and were not fuzzy (Figures 2A and 2B). Upon further examination, we found that AS2^LV^ mutant cells in fact harboured chromosomes that are shorter than those found in wild type cells (Figures 2A and 2C). While the AS1^LV^ mutation leads to hypo-condensation, the AS2^LV^ mutation results in hyper-condensation.

**Figure 2.**
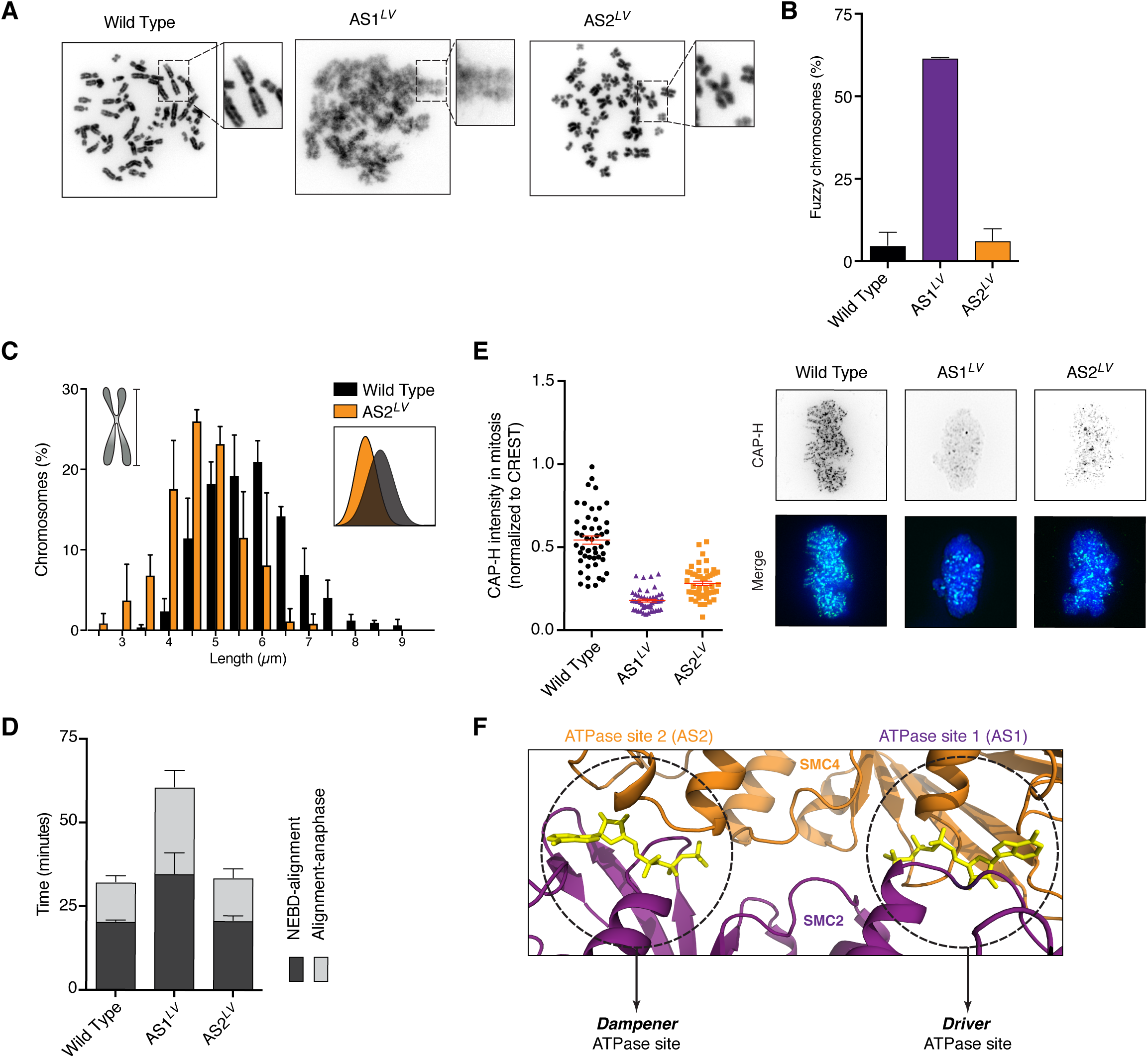
Condensin’s ATPase Sites Drive and Dampen Chromosome Condensation. (A) Representative examples of chromosome spreads of wild type, AS1^LV^ and AS2^LV^ mutant cells. Cells were treated with nocodazole for 1.5 hours prior to shake-off, fixed and stained with DAPI. (B) Percentage of mitotic spreads that display fuzzy chromosomes of cells shown in (A). Error bars represent the SD of three independent experiments. (C) Chromosome length of wild type and AS2^LV^ mutant cells. Chromosomes I, II and III were measured as in the drawn example. The inset depicts a histogram of the chromosome length distribution. Error bars represent the SD of at least three independent experiments. (D) Live-cell imaging of wild type, AS1^LV^ and AS2^LV^ mutant cells. Graph depicts the time spent from nuclear envelope breakdown (NEBD) to alignment and from alignment to anaphase. Cells were transfected with a DNA probe (sirDNA) two hours prior to imaging. Error bars represent the SD of three independent experiments of at least 30 cells per experiment. (E) Quantitative immunofluorescence of chromatin-bound condensin I in wild type, AS1^LV^ and AS2^LV^ mutant cells. Cells were pre-extracted to remove all unbound condensin and then labelled with CAP-H antibody. Intensity of CAP-H is measured and normalised over CREST signal. Each dot depicts the signal of one cell. The red line indicates the mean ± SEM of at least 50 cells per condition. Representative images of mitotic cells stained with CAP-H (green), CREST (cyan) and DAPI (blue) are shown for each genotype. (F) Model for the functional asymmetry of condensin’s ATPase sites. Condensin has two ATPase sites that structurally look symmetrical, but they have distinct functions in the condensation process. We propose that AS1 drives condensation, whereas AS2 acts as a dampener.

The hypo-condensation phenotype can be explained by presuming that condensation requires ATPase activity. But how can we then explain the hyper-condensation phenotype caused by the AS2^LV^ mutation? Hyper-condensed mitotic chromosomes often are a consequence of prolonged mitosis. We therefore followed wild type and mutant cells through mitosis in real-time (Figure 2D). AS1^LV^ cells took longer than wild type to progress from nuclear envelope breakdown (NEBD) until chromosome alignment in metaphase, and these cells also spent more time between alignment and anaphase onset. Presumably both delays are a consequence of the condensation defects observed in these cells. The AS2^LV^ cells, however, progressed normally through mitosis. The observed hyper-condensation effect of the AS2^LV^ mutation therefore is not caused by differences in mitotic timing.

We then measured the levels of the chromatin-bound fraction of condensin in wild type and mutant cells (Figure 2E). Both mutations reduced condensin levels on DNA, which could indicate that both ATPase sites play a role in the loading of condensin complexes onto DNA. To assess the consequences of merely reduced condensin levels, we knocked out one *SMC2* allele in a diploid background. This heterozygous deletion led to a reduction in chromatin-bound condensin that was similar to that observed in AS2^LV^ cells, but resulted in a condensation defect (Figure S2A and S2B). The hyper-condensation therefore is not caused by reduced levels of condensin-bound chromatin.

Cohesin’s release from chromatin is dependent on its release factor WAPL, but also on specifically one of cohesin’s ATPase sites (Beckouët et al., 2016; Çamdere et al., 2015; Elbatsh et al., 2016; Murayama and Uhlmann, 2015). We therefore studied the effects of the AS1^LV^ and AS2^LV^ mutations on the turnover of condensin I on chromatin by fluorescence recovery after photobleaching (FRAP) experiments (Figure S3A and S3B). We could fully recapitulate earlier work with our FRAP assays (Gerlich et al., 2006), but surprisingly we found no evident effect of either AS1^LV^ or AS2^LV^ mutation on condensin’s turnover. This result is important for a number of reasons. First, it indicates that the hypo-and hyper-condensation phenotypes are not caused by effects on condensin’s residence-time on chromatin. Secondly, it indicates that condensin’s DNA release reaction is regulated differently than cohesin’s. This difference may be related to the absence of a WAPL-like release factor for the condensin complex.

The AS2^LV^ hyper-condensation phenotype therefore is not caused by boosting its ATPase activity, nor by affecting mitotic timing. This phenotype also is not caused by affecting condensin’s levels on chromatin, nor by affecting condensin’s residence time on DNA. Together this would indicate that the AS2 ATPase site has a dampening role in the condensation process. The AS2^LV^ mutation results in the functional hyper-activation of condensin, which allows this complex to better condense chromosomes within a given timeframe.

Taken together, our results pinpoint a critical role of condensin’s ATPase in chromosome condensation. There apparently is an asymmetric division of tasks between the two ATPase sites (Figure 2F). One ATPase site (AS1) drives mitotic chromosome condensation, whilst the other ATPase site (AS2) dampens this process. The balanced activity of the driver and dampener ATPase sites appears to be key to achieving wild-type like chromosome condensation.

### Hyperactive Condensin I Shortens Chromosomes in the Total Absence of Condensin II

Metazoans express two distinct condensin complexes. The difference in the localization and stability between condensin I and condensin II hints at specific roles for each of the two complexes. To functionally dissect these specific roles, we knocked out the condensin II specific subunit CAP-H2 in HAP1 cells using CRISPR/Cas9 (Figure 3A). *ΔCAP-H2* cells displayed condensation defects that ranged in severity from cells with long and zig-zag shaped chromosomes to others with an indiscernible fuzzy chromatin mass (Figure 3B and 3E). *ΔCAPH-2* cells also displayed chromosome segregation errors and were delayed in mitosis (Figure 3C, 4A and 4B).

**Figure 3.**
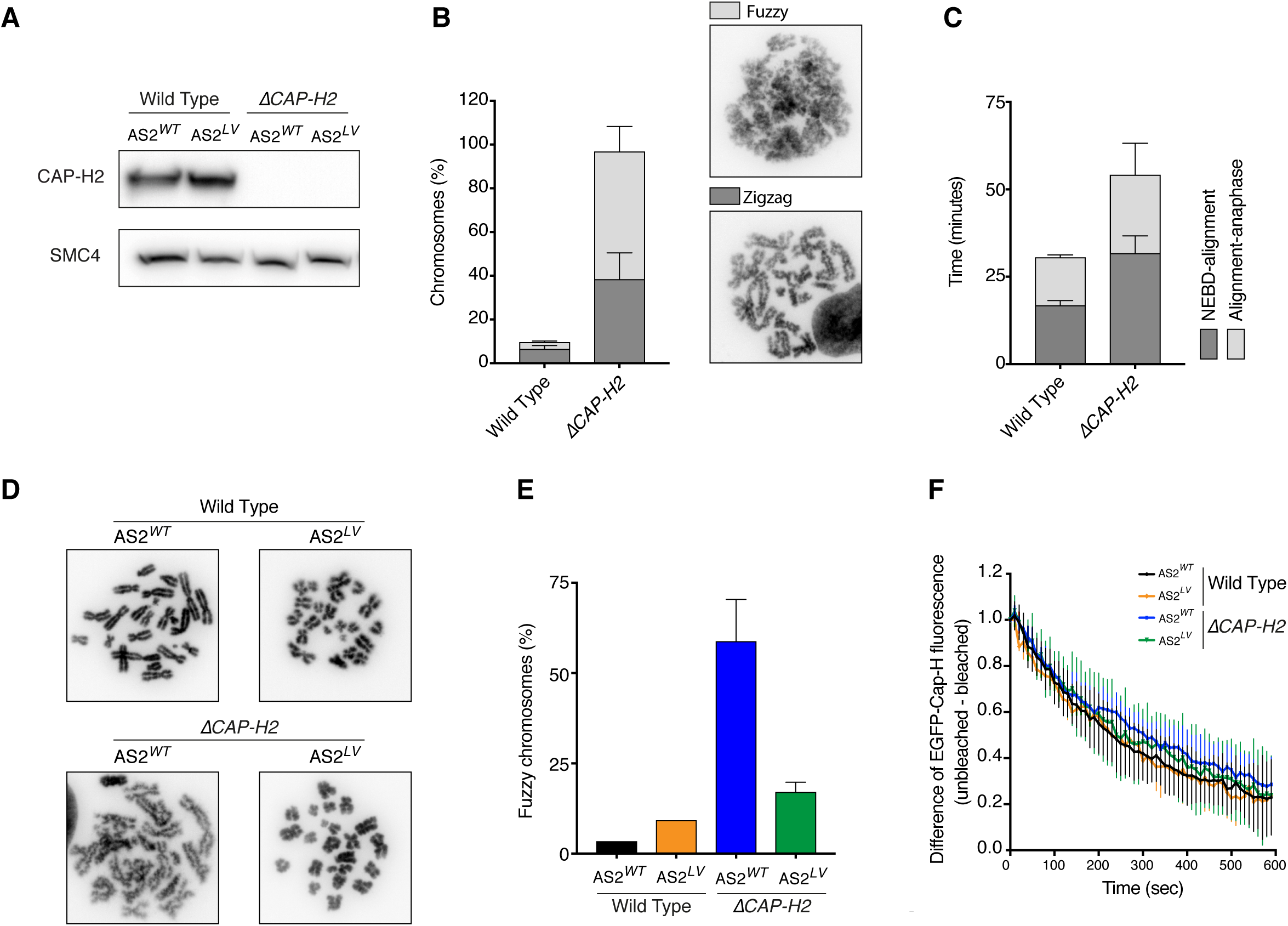
Hyperactive Condensin I Condenses Chromosomes in the Total Absence of Condensin II. (A) Western blot analyses of CAP-H2 in wild type and *ΔCAP-H2* cells with either AS2^WT^ or AS2^LV^. SMC4 is used as a loading control. (B) Percentage of mitotic spreads with either fuzzy or zigzag-shaped chromosome morphology in wild type and *ΔCAP-H2* cells. Error bars represent the SD of three independent experiments of at least 120 chromosome spreads per experiment. Representative images are shown. (C) Live-cell imaging of wild type and *ΔCAP-H2* cells as shown in Figure 2E. (D) Representative images of chromosome spreads of wild type and *ΔCAP-H2* cells with either AS2^WT^ or AS2^LV^. At least 120 spreads were examined from two independent experiments. (E) Percentage of mitotic spreads with fuzzy chromosome morphology for the indicated genotypes. Error bars represent the SD of three independent experiments of at least 120 chromosome spreads for each experiment. (F) FRAP experiments in the indicated genotypes. Error bars represent SD of three different experiments, except for wild type AS^LV^, which is from one experiment.

**Figure 4.**
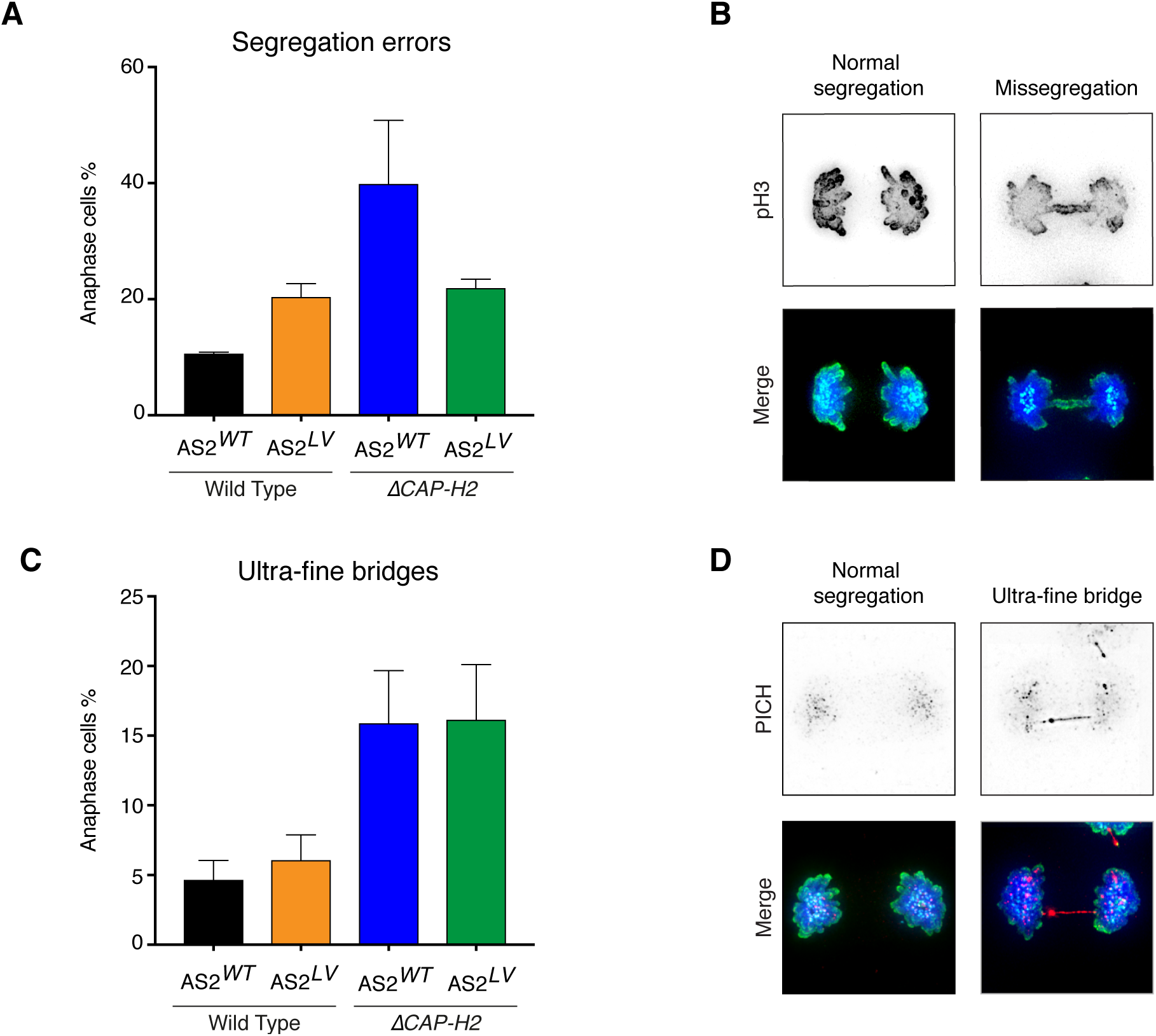
Condensin II Has a Specific Role in Sister Chromatid Decatenation. (A) Quantification of segregation errors that occurs during anaphase for the indicated genotypes. DAPI (blue), phospho-histone H3 (pH3) (green) and CREST (cyan) stainings were visualized by immunofluorescence microscopy. Error bars represent the SD of two independent experiments of at least 50 anaphases for each experiment. (B) Representative examples of an anaphase with either normal segregation or a missegregation that are quantified in (A). (C) Quantification of ultra-fine bridges (UFBs) visualised by PICH stainings. DAPI (blue), and CREST (cyan) and PICH (red) stainings were visualized by immunofluorescence microscopy. Error bars represent the SD of three independent experiments of at least 50 anaphases for each experiment. (D) Representative examples of an anaphase with either normal segregation or a segregation with an UFB that are quantified in (C).

We then examined which *ΔCAP-H2* defects can or cannot be rescued by boosting the activity of condensin I with our hyper-activating AS2^LV^ mutation. We therefore introduced this mutation into *ΔCAP-H2* HAP1 cells. Strikingly, chromosomes of *ΔCAP-H2* cells expressing the AS2^LV^ mutant condensed well, and generally were even shorter than wild type chromosomes (Figure 3D). The AS2^LV^ mutation also rescued the high percentage of fuzzy chromosomes caused by condensin II deficiency (Figure 3E). Notably, condensin I turnover was not affected by condensin II deficiency, nor by the AS2^LV^ mutation (Figure 3F). Condensin II, with its stable DNA-binding mode therefore is neither inherently required for efficient condensation, nor for the shortening of chromosomes in particular.

### Condensin II Has a Specific Role in Sister Chromatid Decatenation

Proper condensation is a prerequisite for faithful chromosome segregation (Gerlich et al., 2006; Nagasaka et al., 2016). Consistently, *ΔCAP-H2* cells suffer from an increase in missegregations compared to wild type cells (Figure 4A and 4B). In correspondence with the rescue of the major condensation defect, the hyper-activating AS2^LV^ mutant could also largely rescue this segregation defect. Condensin II deficiency did not affect basic chromosome segregation in AS2^LV^ mutant cells. However, cells harbouring this mutation did not reach wild type segregation fidelity, indicating that hyper-condensation may not be beneficial for faithful chromosome segregation.

Mitoses of *ΔCAP-H2* cells also displayed an increase in ultrafine bridges (Figure 4C and 4D). This is interesting, as condensin II is nuclear throughout the cell cycle, and it plays a role in the resolution of sister chromatids from S phase onwards (Ono et al., 2013). This resolution could for example aid the disentangling or decatenation of sister DNAs, and the persistence of such catenanes can lead to ultrafine DNA bridges in mitosis. Condensin I only contacts DNA upon nuclear envelope breakdown (NEBD). Importantly, we find that hyperactive condensin I cannot rescue the increase in ultrafine bridges observed in *ΔCAP-H2* cells. Together, this would support the model that hyperactive condensin I by itself can condense chromosomes within the relatively short window spanning from NEBD to metaphase, but that decatenation requires more time, and therefore needs condensin II from S phase onwards.

### Condensin II Has a Specific Role in Mitotic chromosome organization

Upon closer examination of the hyper-condensed chromosomes, we noticed that chromosomes of *ΔCAP-H2* cells harbouring the hyper-activating AS2^LV^ mutant appeared less straight than those of AS2^LV^ single mutant cells. We therefore further examined mitotic chromosome organization by performing Hi-C (Figure 5A). Wild type cells, in correspondence with earlier work (Naumova et al., 2013), yielded a Hi-C map with homogenous interactions along the whole length of chromosomes. This uniform interaction pattern yielded a band enriched for *cis* contacts up to ~10Mb distance away from the main diagonal. AS2^LV^ mutant cells interestingly displayed a band with longer-distance contacts that now increased to ~25Mb (Figure 5A). These findings are supported by relative contact probability plots (Figure 5B). The plots of wild type cells display an increase in contacts at ~10Mb distance, which shifts to ~25Mb for the AS2^LV^ mutant cells. We suggest that this increase in contact length is caused by the formation of extended chromatin loops by hyperactive AS2^LV^ condensin.

**Figure 5.**
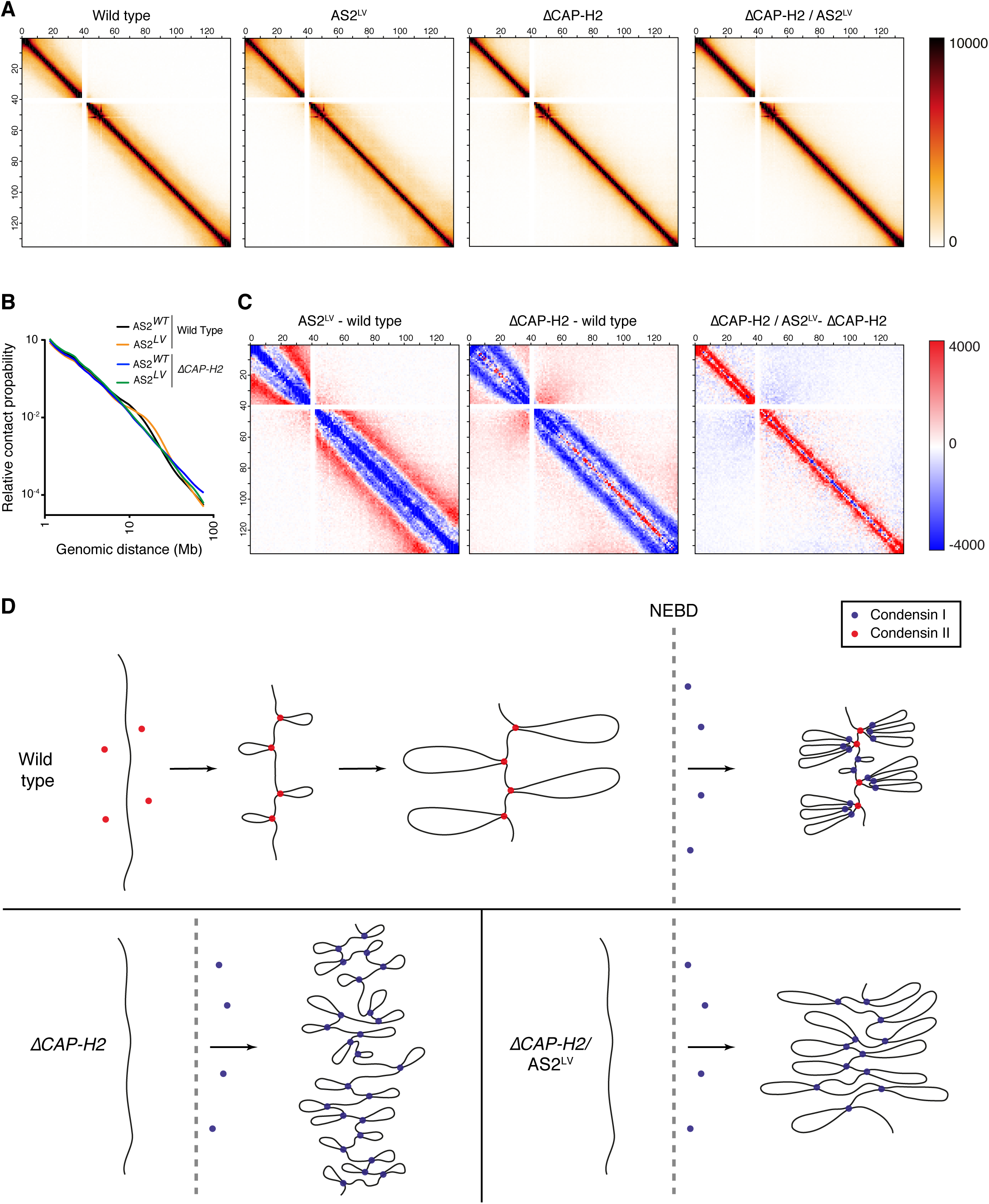
Condensin II Has a Specific Role in Mitotic Chromosome Organization. (A) Whole-chomosome Hi-C matrices of mitotic cells. Chromosome 10 is shown at 1 Mb resolution. Cells were harvested in mitosis by mitotic shake-off after 1.5 hours Nocodazole treatment. The harvested cells were further selected for their mitotic status by FACS-sorting for phospho-MPM2 positivity. (B) Relative contact probability plot displaying the likelihood of a contact at increasing length scales of the cells as shown in (A). (C) Differential Hi-C signal between the indicated cell lines. (D) A tentative model for the specific roles of condensin I and II complexes in mitotic chromosome condensation.

Strikingly, neither *ΔCAP-H2* cells, nor *ΔCAP-H2* cells with the AS2^LV^ mutation yielded a band with contacts over ~5Mb distance. Such a condensin II dependent role in the formation of longer-range contacts is in correspondence with a recent preprint using mitotic chicken cells (Gibcus et al., 2017). How then can the AS2^LV^ mutant yield hyper-condensed chromosomes both in wild type and in *ΔCAP-H2* cells, if their interaction maps look so different? We therefore plotted the difference in interactions between the respective genotypes (Figure 5C). We found that AS2^LV^ mutant cells displayed longer-distance contacts up to ~25 Mb relative to wild type. The AS2^LV^ mutant in *ΔCAP-H2* cells in fact also displayed an increase in contact frequency, but here they shifted from very short-range contacts close to the diagonal of the Hi-C map to now ~5 Mb contacts.

Our results therefore indicate that condensin I and II each form loops with specific ranges in size. Condensin II forms long loops, while condensin I forms smaller loops. Hyper-activation of either of these complexes by the AS2^LV^ mutation appears to yield extended loops within the size-range of the respective complex. Even though the AS2^LV^ mutant yields hyper-condensed chromosomes both in wild type and in *ΔCAP-H2* cells, and chromosomes superficially look similar, condensin II does have a specific role in the organization of these chromosomes. We propose that the formation of condensin II-specific long loops is required for a straight chromosomal axis (Figure 5D).

## DISCUSSION

### Condensin’s Driver and Dampener ATPase Sites

What drives mitotic chromosome condensation is a major question in chromosome biology. We here identify a key role of condensin’s ATPase machinery in this process. Specifically one of condensin’s ATPase sites turns out to drive condensation. The other ATPase site however dampens this process. The current model is that condensin promotes condensation through the processive enlargement of chromatin loops (Goloborodko et al., 2016; Nasmyth, 2001). Our data suggests that condensin’s AS1 ATPase site indeed provides the driving force behind this loop enlargement.

How then does the dampener ATPase site affect loop formation? We find that mutating this AS2 site yields more contacts over longer distances as compared to wild type. The AS2^LV^ mutant achieves this without affecting mitotic timing. This is important to know, as prolonged mitosis is a common cause for hyper-condensation. It does not affect condensin’s turnover on chromatin either. This would have been a valid explanation for the observed hyper-condensation, as stabilising cohesin on chromatin leads to longer loops in interphase chromosomes (Haarhuis et al., 2017). If condensin’s AS2^LV^ mutation affects neither mitotic timing nor condensin’s stability, this would leave an increase in condensin’s speed as a likely explanation for the phenotype. A faster, hyperactive, condensin could then achieve hyper-condensation within a normal time-frame.

Recent work on bacterial SMC indicates that this complex can exist in either a closed or an open conformation (Diebold-Durand et al., 2017), and that the ATPase machinery controls the switch between these two states. This raises the interesting possibility that a repetitive switch between these two states drives loop enlargement. Condensin’s driver ATPase site could trigger this switch and thus lead to loop enlargement. The dampener site on the other hand could antagonize this switch by keeping the complex in an intermediate, non-productive, state.

An asymmetric division of tasks may well reflect a universal principle for SMC ATPases. Cohesin’s ATPase machinery for example also harbours an asymmetric activity, as specifically one of cohesin’s ATPase sites is required for its release from DNA (Beckouët et al., 2016; Çamdere et al., 2015; Elbatsh et al., 2016; Murayama and Uhlmann, 2015). Our FRAP experiments however indicate that the analogous ATPase site in condensin (AS2) does not play such a role. This may well be related to the fact that condensin has no dedicated WAPL-like release factor. How condensin in fact releases DNA remains a mystery. It is striking though that this same site in both cases has a function that is antagonistic to the overall function of the complex. For condensin, this site has a dampening function, while for cohesin it is required for the release of this complex from chromatin. Condensin’s HEAT repeat subunits have contrasting functions in condensation (Kinoshita et al., 2015). Whether these opposing functions are linked to the ATPase machinery is unknown. Each of these proteins binds to a different part of condensin’s kleisin subunit (Piazza et al., 2014). Recent work on bacterial SMCs indicates that the individual ATPase sites can be regulated independently through binding to different parts of the kleisin (Zawadzka et al., 2017).

Recent single-molecule experiments indicate that condensin is a molecular motor that can translocate along DNA in an ATP-dependent manner (Terekawa et al., 2017), and that ATP hydrolysis is required both for stable, salt-resistant DNA binding and for subsequent compaction of DNA (Eeftens et al., 2017; Strick et al., 2004). Hydrolysis-deficient condensin complexes were previously found at chromatin (Hudson et al., 2008; Kinoshita et al., 2015), but considering this in vitro data, these complexes presumably were not stably DNA bound. This is in line with experiments using these classical ‘Walker B’ mutant forms of the cohesin complex (Hu et al., 2011; Ladurner et al., 2014).

The AS1^LV^ and AS2^LV^ mutants are partially impaired in their ATPase activity, but they do bind DNA with wild type-like stability. It could be argued that these mutants reflect hypo-morphs of sorts. This partial inactivation however has allowed us to also investigate the roles of the different ATPase sites post condensin loading. Our data would be in line with the single-molecule experiments, but the loading and condensation processes do appear to have different requirements for each ATPase site. We show that both sites are involved in loading, but that specifically AS1 is important for the condensation process itself, which in turn is dampened by AS2.

### Condensin I Versus Condensin II

Metazoans express two distinct condensin complexes that differ in their protein composition, cellular localization, and stability on DNA. We find that condensin II deficient cells have long and fuzzy chromosomes, but that hyperactive condensin I can efficiently shorten chromosomes even in the total absence of condensin II. This is surprising, because condensin II is thought to have a specific role in the shortening of mitotic chromosomes (Green et al., 2012; Shintomi and Hirano, 2011). We propose that the time from NEBD to anaphase is not sufficient for wild type condensin I to fully condense chromosomes without the help of condensin II, but that condensin II does not have an irreplaceable function in this process.

We find that condensin II does have at least two key roles that cannot be compensated for by hyperactive condensin I. The first is the decatenation of sister chromatids. The nuclear localization of condensin II allows this complex to aid decatenation from S phase onwards. Even though hyperactive condensin I is sufficient for the condensation process per se within the limited time-window from NEBD to anaphase, the decatenation process may well require considerably more time and mostly takes place during interphase. Condensin I can in fact regulate DNA catenation, as its inactivation in *Drosophila* cells during metaphase results in an increase in catenanes (Piskadlo et al., 2017). We therefore expect that condensin II’s contribution to decatenation simply is a consequence of the spatiotemporal regulation of this complex.

Second, we show that condensin II has a specific role in the formation of longer-range *cis* contacts than does condensin I. Considering these findings, and the timing at which the two complexes contact DNA, our data would support the model that condensin II initiates condensation by forming long loops, and that upon NEBD, condensin I makes small loops within these larger loops (Figure 5D). This fits also with earlier work showing that condensin I is important for reducing the width of chromosomes (Green et al., 2012; Shintomi and Hirano, 2011). Condensin II in a normal setting would then be important for the shortening of chromosomes, but we find that this role can be taken over by hyperactive condensin I. Importantly though, these shortened chromosomes are morphologically distinct, as they appear less straight. The condensin II-specific long loops may be important for the formation of straight chromosomes.

A recent preprint suggests that condensin II plays a role in the formation of a helical turn within the axis of mitotic chromosomes, and that megabase-scale contacts reflect interactions between loops that are stacked in a helical fashion around this axis (Gibcus et al., 2017). Our data would be in line with this hypothesis. We can however also explain our data by presuming that the bulk of these contacts reflect DNA-DNA interactions at the base of loops. Then condensin II would normally produce loops that extend up to ~10 Mb, and condensin I would produce shorter loops within and outside the condensin II loops. In this latter scenario, the antler-shaped structures of loops within loops would stack on top of each other in the energetically most favoured manner, without the requirement for a helical axis. The formation of short loops within longer ones would then be key to achieving a straight, rigid, chromosomal axis.

Hyper-activation of both condensin I and condensin II leads to hyper-condensed chromosomes, but interestingly this also causes an increase in chromosome missegregations. These results imply that condensin’s ability to condense chromosomes in fact needs dampening to ensure faithful chromosome segregation. The balanced activity of the driver and dampener ATPase sites indeed appears to be key to both the condensation and the segregation of chromosomes. Why the degree or speed of condensation needs to be kept in check is an important question for the future.

## ACKNOWLEDGEMENTS

We are grateful to all members of the Rowland and Medema Laboratories for helpful discussions, Roy van Heesbeen and Alberto García Nieto for technical assistance, Bram van den Broek for advice on imaging, and the NKI protein production, Microscopy, and FACS facilities for assistance. We thank Kim Nasmyth for discussions and Martin Houlard for imaging in murine cells. We acknowledge the Netherlands Organisation for Scientific Research (ECHO) for funding.

## METHODS

### Cell culture and transfections

HAP1 cells were cultured at 37°C with 5% CO_2_ in IMDM (Gibco), supplemented with 12% FCS (Clontech), 1% Penicillin/Streptomycin (Invitrogen) and 1% UltraGlutamine (Lonza). DNA transfections were performed using FuGENE 6 (Promega) according to the manufacturer’s protocol.

### Genome editing

To introduce endogenous mutations, CRISPR/Cas9 genome-editing was performed as described (Elbatsh et al., 2016). In brief, guide RNAs (gRNAs) targeting SMC2 and SMC4 were designed using the online CRISPR design tool (crispr.mit.edu). SMC2 gRNA sequences used: Forward CACCGCTGAACTTAGTGGTGGTCAG, Reverse AAACCTGACCACCACTAAGTTCAGC. SMC4 gRNA sequences used: Forward CACCGAAGATCTTCAACCTTTCGGG and Reverse AAACCCCGAAAGGTTGAAGATCTTC. Phosphorylated and annealed oligo’s were ligated into pX330 (Addgene plasmid #42230). To introduce the desired mutations we designed a 90-bp homology-directed donor oligo: for SMC2: CAAGGTTGCCTTGGGAAATACCTGGAAAGAAAACCTAACTGAAGTTAGTGGTGGTCAGAGGTGAGGAATCACTTTGCTATATTATAATTT. For SMC4’s donor oligo: CAGTGTTCGACCACCTAAGAAAAGTTGGAAAAAGATCTTCAACGTTTCGGGAGGAGAGAAAACACTTAGTTCATTGGCTTTAGTATTTGC. To obtain *CAP-H2* knockout cells, Forward CACCGTTCTTGGTGAGGTCGCGGA and Reverse AAACTCCGCGACCTCACC AAGAAC sgRNAs were cloned into px330. HAP1 clones were generated by insertion of a Blasticidine cassette, as previously described (Haarhuis et al., 2017). 0.1μg pBabe-Puro was co-transfected (1:10 ratio to CRISPRs) to select for transfected cells by puromycin (2 μg/ml, Sigma-Aldrich). Genomic DNA was isolated from picked clones and the desired mutations were scored for by Sanger Sequencing.

### ATPase assays

The five-subunit wild type or mutant condensin complexes were expressed and purified, and ATP hydrolysis rates of condensin holocomplexes were measured as described (Terekawa et al., 2017). In short, reactions (10μl) were prepared with 0.5 μM condensin holocomplex, with or without 24 nM relaxed circular 6.4-kb plasmid DNA in ATPase buffer (40 mM TRIS-HCl pH 7.5, 125 mM NaCl, 10% (v/v) glycerol, 5 mM MgCl_2_, 5 mM ATP, 1 mM DTT and 33 nM [α^32^P]-ATP; Hartmann Analytic) and incubated at room temperature (~25°C). One microliter of the reaction mix was spotted onto PEI cellulose F TLC plates (Merck) every 3 min for a total duration of 15 min. Reaction products were resolved using 0.5 M LiCl and 1 M formic acid solution and analyzed on a Typhoon FLA 9,500 scanner (GE Healthcare). ATP hydrolysis rates were calculated from the ADP/ATP ratios from time points in the linear range of the reaction.

### Immunofluorescence

Cells were grown on 12 mm glass coverslips and pre-extracted using PEM-T buffer (100mM PIPES pH 6.8, 1 mM MgCl_2_, 5 mM EGTA and 0.2% Triton) for 60 seconds. Fixation was performed with 4% paraformaldehyde for 6 minutes with PEM-T buffer followed by 4 minutes without and blocking was with 3% BSA in PBS for 1 hour. The following antibodies were used: SMC2 (Bethyl, A300-058A), CREST (Cortex Biochem), PICH (Abnova). All antibodies were used at 1:1000 dilution and incubated overnight at 4°C. Secondary antibodies (Molecular probes, Invitrogen) and DAPI were incubated for 1 hour at room temperature. Coverslips were mounted with Vectashield (Vector Laboratories, H-1000). Images were taken using a Deltavision deconvolution microscope (Applied Precision) and image acquisition was done using Softworx (Applied Precision), ImageJ and Photoshop.

### Hi-C

Cells were plated in T175 flasks (Cellstar, Greiner bio-one). The next day, cells were treated with nocodazole for 1.5 hours and mitotic cells were collected by shake-off. Cells were cross-linked with 2% formaldehyde, blocked with 3% BSA in PBST and then incubated with the mitosis-specific antibody MPM2 (1:500, Millipore) overnight at 4°C. Cells were washed with PBST then incubated with the secondary Alexafluor antibody. Labelled mitotic cells were sorted by FACS (BD FASAira II, BD biosciences). 5-10 million sorted cells were treated with 0.1% SDS final concentration (5μl 10%SDS per 500μl volume) and incubated for 10 minutes at 65°C. Then 30μl 20% triton was added to the cells (final concentration= 1% triton). Afterwards, sample preparation was done as described (Haarhuis et al., 2017). Raw sequence data was mapped and processed using HiC-Pro v2.9 (Servant et al., 2015). Data was mapped to hg19. The relative contact probability was computed as previously described (Haarhuis et al., 2017).

### Chromosome spreads

Cells were treated with nocodazole for 1.5 hours and mitotic cells were collected by shake-off. Cells were incubated with 75 mM KCl for 20 minutes. Cell pellets were twice resuspended in fixative solution (Methanol: Acetic Acid, 3:1) and spun down. Fixed cells were incubated with DAPI and dropped on tilted cover slides, and then dried at 37°C. Dried slides were mounted with Vectashield (Vector Laboratories, H-1000). Images of spreads were taken using the Metafer system (Metasystems).

### Live-cell imaging

Cells were grown on 8-well glass-bottom dishes (LabTek). Two hours prior to imaging, sirDNA probe (1:2000, Spirochrome) was added together with verapamil (1:10000) to visualize DNA in L-15 CO_2_-independent medium (Gibco). Images were taken using a Deltavision deconvolution microscope (Applied Precision). Cells were imaged every 5 minutes using a 40x air objective with 3 × 3 μm Z-stacks. Image acquisition was carried out using Softworx (Applied Precision) and ImageJ.

### Fluorescence Recovery after Photobleaching (FRAP)

Cells were transfected with a pC1-EGFP-CAP-H construct in 4-well glass bottom dishes (LabTek). 48 hours post-transfection, sirDNA probe (1:2000, Spirochrome) was added to cells to visualize DNA, in L-15 CO_2_-independent medium (Gibco). FRAP experiments were performed using a confocal microscope (Leica Microsystems) with a 63x oil immersion objective using the LAS-AF FRAP-Wizard. Images were taken by accumulating 4 frames at a scanning speed of 1000 Hz. Photobleaching was performed by 6 times illuminating the selected region at 100-fold laser power (488 nm Laser). 60 post-bleaching images were taken with 10 second intervals. Analysis of images was done using an in-house ImageJ macro. A mask on the DNA signal was created and the EGFP-CAP-H signal was normalized to the DNA signal. Graphpad Prism 6 was used to fit the data using a one phase decay function.

### Western Blotting

Cells were grown in 6-well plates, harvested and lysed with Laemmli buffer (120 mM Tris pH 6.8, 4% SDS, 20% glycerol) by boiling at 95°C for 5 minutes. Total protein concentration was determined with Lowry assays and protein levels were analysed with western blotting. The following antibodies were used: SMC4 (1:5000 Bethyl A300-064a) and CAP-H2 (1:1000, Bethyl, A302-275a).

---

**Figure S1.**
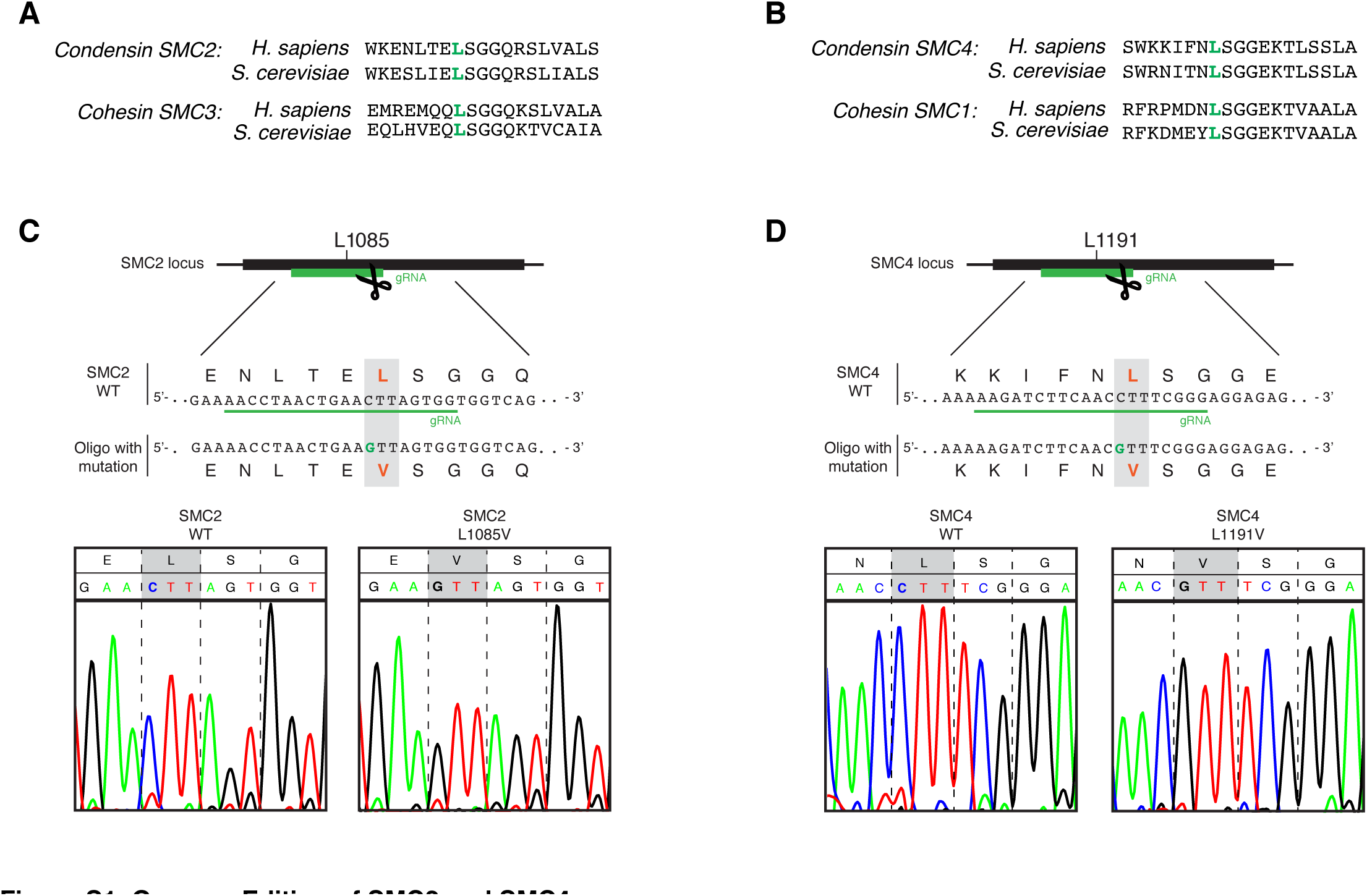
Genome Editing of SMC2 and SMC4. (A) Sequence alignment showing the conservation of a key amino acid in the signature motif of the ATPase domains of conden-sin’s SMC2 and cohesin’s SMC3. (B) Sequence alignment showing the conservation of a key amino acid in the signature motif of the ATPase domains of conden-sin’s SMC4 and cohesin’s SMC1. (C) CRISPR/Cas9 genome-editing technology used to make the L1085V mutation in SMC2 in human HAP1 cells. The guide RNA (gRNA) was designed to target Cas9 to cut in the signature motif of the SMC2 gene and the L1085V (C>G) mutation was introduced through homology-directed repair using a 90 bp donor-oligo that carries the mutation. Chromatograms of the Sanger sequencing of cells with SMC2 wild type (left) and SMC2 L1085V (right) are shown. (D) Same as in (C) but for the SMC4 L1191V mutation.

**Figure S2.**
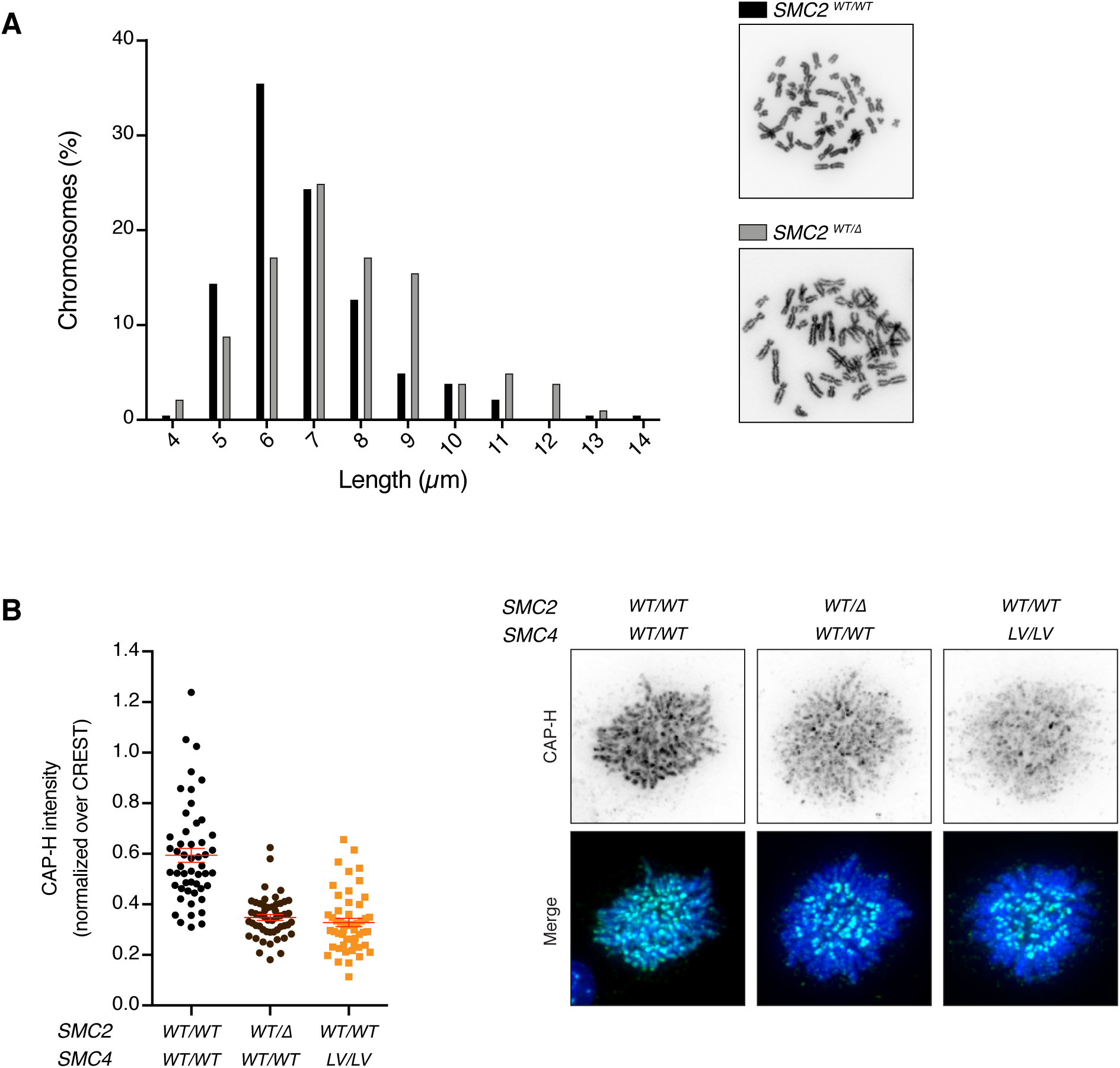
Reducing Condensin Levels on Chromatin Impairs Mitotic Chromosome Condensation. (A) Chromosome length of wild type and *SMC2*^*WT*/∆^ mutant cells. Diploid derivatives of HAP1 cells were transfected with a guide RNA targeting Cas9 to the *SMC2* gene. Chromosome spreads were prepared of cells harbouring a heterozygous deletion of *SMC2* and of a diploid wild type control. (B) Quantitative immunofluorescence of chromatin-bound condensin I in wild type, *SMC2*^*WT/∆*^ and *SMC4^LV/LV^* (AS2^LV^) mutant cells. Cells were pre-extracted to remove all unbound condensin and then stained with a CAP-H antibody. Intensity of CAP-H was measured and normalised over CREST signal. Each dot depicts the signal of one cell. The red line indicates the mean ± SEM and n is at least 50 cells per condition. Representative images of mitotic cells stained with CAP-H (green), CREST (cyan) and DAPI (blue) are shown for each genotype.

**Figure S3.**
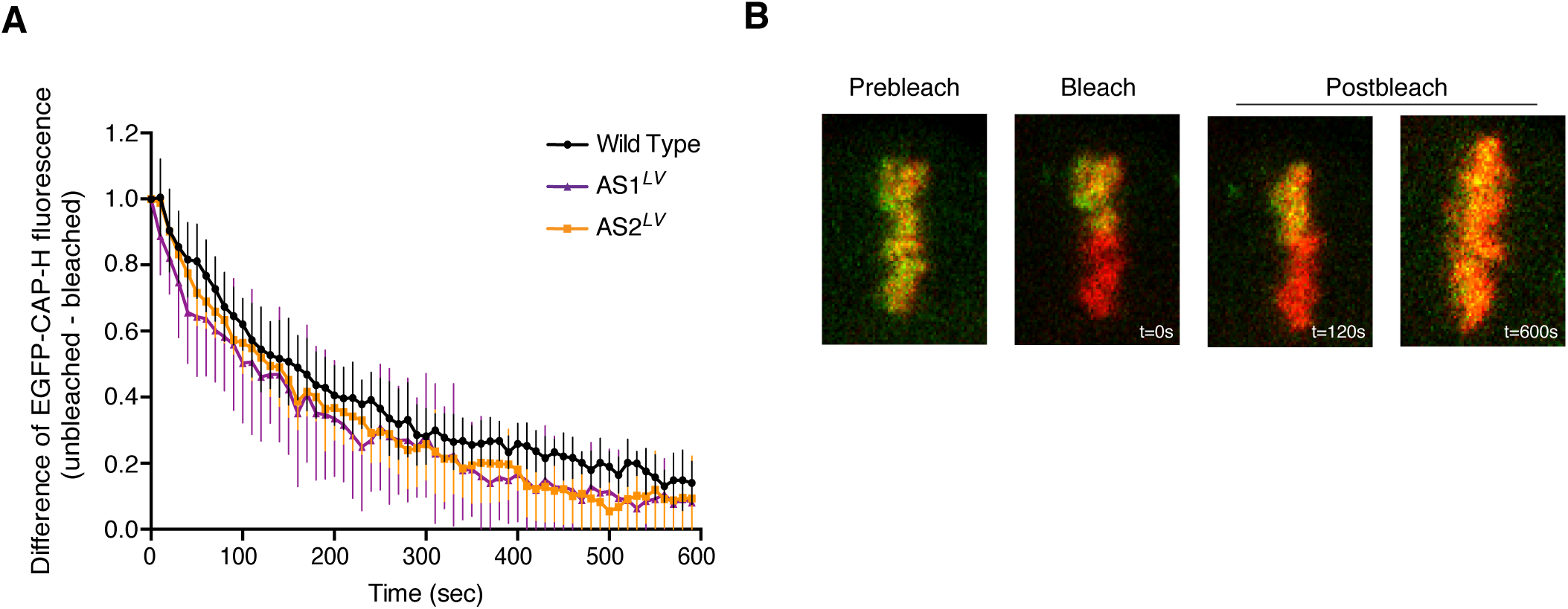
AS2^LV^ and AS2^LV^ ATPase Mutants Do Not Affect Condensin I’s Turnover in Mitosis. (A) Wild type, AS1^LV^ and AS2^LV^ mutant cells expressing EGFP-CAP-H were used for fluorescence recovery after photobleaching (FRAP) experiments. The entire fluorescent signal in cells was bleached except for half of the metaphase plate. Recovery in the bleached and unbleached halves of the plate was followed by time-lapse imaging. DNA was visualized using a DNA probe (sirDNA). The curves represent the difference in fluorescence signal between bleached and unbleached regions after normalization with the DNA signal. n= ≥ 7 cells from at least two independent experiments. Errors bars represent the SEM. (B) Representative images before, at, or after bleaching for the cells in (A) are shown with the indicated time in seconds.

